# Homologues of the inner-membrane LPS transport proteins are required for sphingolipid transport in *Caulobacter crescentus*

**DOI:** 10.64898/2026.04.12.717747

**Authors:** Chioma G. Uchendu, Georgia L. Isom, Eric A. Klein

## Abstract

Recent elucidation of the bacterial sphingolipid synthesis pathway has revealed that these lipids are produced by a range of taxonomically diverse species. In contrast to the biosynthetic pathways, the mechanism by which sphingolipids are transported from the inner membrane to the cell surface in Gram-negative bacteria remains a mystery. Here, we identify and characterize paralogs of the well-characterized lipopolysaccharide (LPS) inner membrane ABC transporter proteins encoded within the sphingolipid locus. Using *Caulobacter crescentus* as a model system, we analyzed three putative inner membrane proteins with homology to LptF, LptG, and LptC. Deletion of these genes was lethal, likely due to the accumulation of anionic sphingolipids in the inner membrane. We further show that the LptF and LptG homologues form a complex like their LPS counterparts and discover that they interact with the LPS ATPase LptB. Together, our data suggest that ceramide transport to the outer membrane is facilitated by an ABC transporter consisting of a sphingolipid-specific LptFG homolog coupled to the LPS LptB, supporting a model in which sphingolipid transport partially converges with the LPS transport system. Together, these findings reveal an unexpected evolutionary relationship between sphingolipid and lipopolysaccharide transport.

## Introduction

Gram-negative bacteria have two structurally and functionally distinct membranes: the inner membrane (IM) and the outer membrane (OM). The IM encapsulates the cytoplasmic contents and is separated from the OM by an aqueous compartment known as the periplasm (1) (2). The IM is primarily composed of phospholipids and integral transmembrane (TM) proteins that span the bilayer through alpha-helical transmembrane domains (3).

By contrast, the OM exhibits asymmetric lipid organization; phospholipids make up the inner leaflet, while lipopolysaccharide (LPS) constitutes the majority of the outer leaflet (4). Embedded within the OM are the integral outer membrane proteins (OMPs), which cut across the membrane as antiparallel beta strands that fold into beta-barrel structures (5). This asymmetric structure creates a permeability barrier, preventing the free flow of large molecules and limiting the entry of small hydrophobic compounds into the cell (2). Thus, the barrier function of the OM is intrinsically linked to its unique lipid composition and structural organization (6).

LPS is the defining constituent of the outer membrane for most Gram-negative bacteria, however notable exceptions exist. Organisms such as the hyperthermophile *Aquifex pyrophilus,* spirochetes *Treponema pallidum* and *Borrelia burgdorferi*, and members of the genus *Sphingomonas* lack canonical LPS in their outer membranes. Instead, these species utilize alternative glycolipids (7) or glycosphingolipids (8).

Even for organisms that routinely produce LPS, these lipids may not always be absolutely essential. To date, LPS-deficient mutants have been isolated for *Moraxella catarrhalis*, *Acinetobacter baumanii*, *Neisseria meningitidis*, and *Caulobacter crescentus* (9–12). In the case of *C. crescentus*, we recently demonstrated that this survival is mediated by the production of a novel anionic sphingolipid, ceramide diphosphoglycerate (CPG2) (12). Colistin sensitivity experiments demonstrated that, in *C. crescentus*, colistin acts by binding CPG2 on the cell surface rather than LPS (12, 13). Furthermore, sphingolipid-deficient *C. crescentus* have increased sensitivity to salt, temperature, and detergents (14, 15). Together, these findings establish that outer membrane sphingolipids have critical physiological roles in membrane and cellular integrity.

While the biochemical pathways for sphingolipid synthesis are fairly well characterized (14–18), nothing is known regarding how these lipids are translocated to the cell surface. The initial stages of sphingolipid biosynthesis occur in the cytoplasm (19); thus, sphingolipids require transit across the inner membrane and periplasm before arriving on the outer leaflet of the OM.

The shuttling of sphingolipids from the IM to the OM outer leaflet is likely reminiscent of LPS trafficking. Transport of LPS begins with the essential ABC transporter MsbA, which facilitates ATP-dependent lipid flipping across the IM to the periplasm (20). The LptB_2_FGC complex then extracts LPS from the IM and transfers it to LptA which forms a bridge across the periplasm. Finally, LPS is delivered to LptD/E at the OM for outer leaflet insertion. (21).

In this report, we characterize homologues of LptFGC encoded in the sphingolipid locus of *C. crescentus*. These genes are conserved in many sphingolipid-producing bacteria and their essentiality under conditions of sphingolipid synthesis implicates them in sphingolipid transport. Affinity co-purification assays demonstrate that the LptF and LptG homologues interact with each other, as well as with the LPS-associated IM ATPase LptB. Our findings reveal a new inner membrane lipid transport complex implicated in sphingolipid OM trafficking.

## Results

### Sphingolipids are required for optimal outer-membrane function

Although sphingolipids are dispensable for growth under standard laboratory conditions (13), loss of these lipids increases sensitivity to salt and detergent stress (15). In the absence of sphingolipids (Δ*spt*), the outer membrane (OM) became more sensitive to agar plates containing the antibiotics bacitracin and vancomycin, which generally do not efficiently pass through the OM of Gram-negative bacteria (Figure 1A). Furthermore, the Δ*spt* strain is more permeable to fluorescent vancomycin (Figure 1B). Similarly, the lack of sphingolipids sensitizes the bacteria to bile salts such as deoxycholate (15) (Figure 1C). All of these phenotypes could be rescued by ectopic expression of *spt* from a vanillate-inducible promoter (Figures 1A-C).

**Figure 1:**
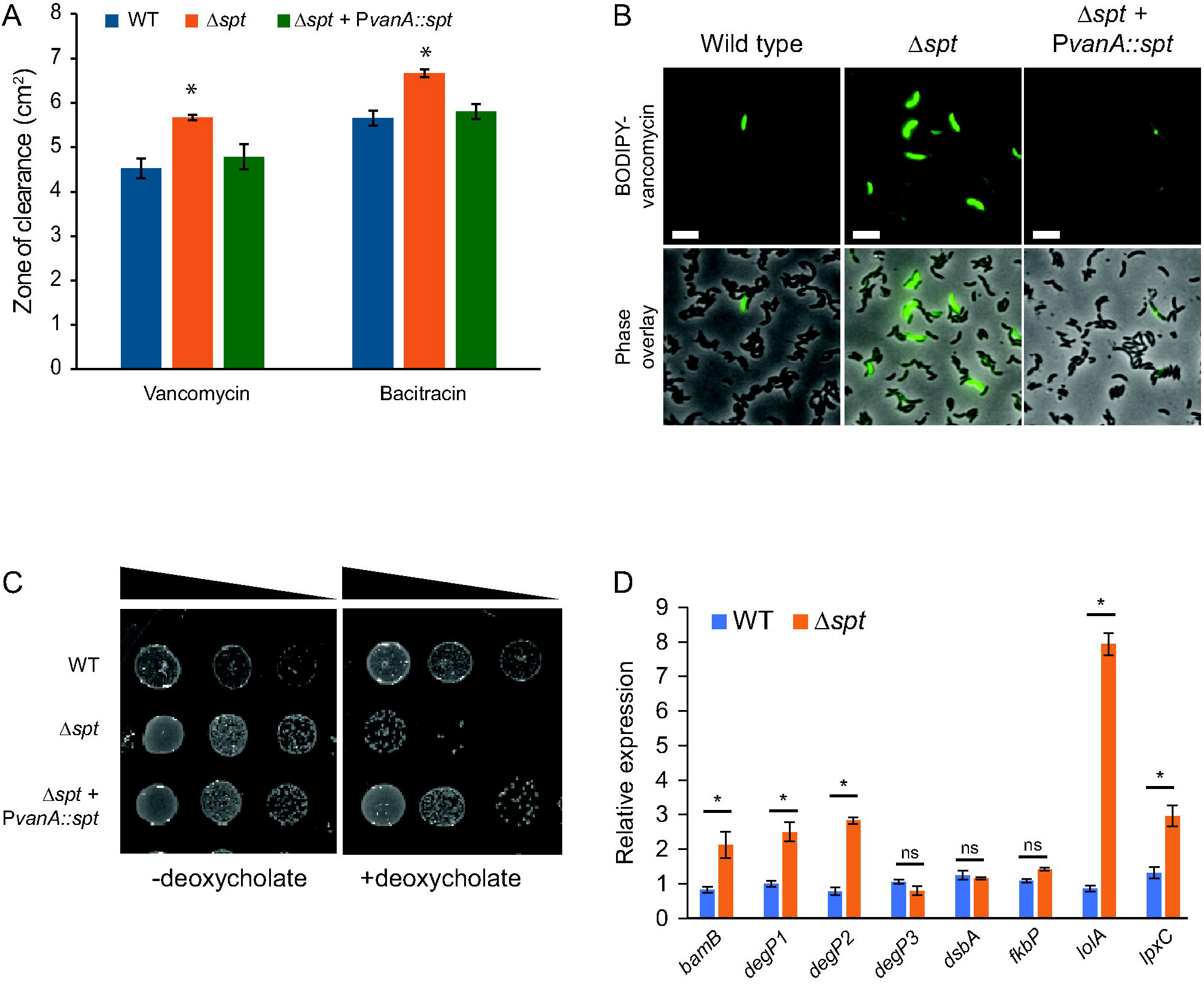
Sphingolipids contribute to outer membrane homeostasis. (A) Antibiotic disc-diffusion assays were performed on the indicated strains. Δ*spt* cells cannot produce any sphingolipids. The complementation strain (Δ*spt* + P*vanA*::*spt*) was induced with 0.5 mM vanillate. The disks contained either vancomycin (0.4 mg) or bacitracin (0.2 mg). The area of the zone-of-clearing was measured from the center of the filter disc. An analysis of variance (ANOVA) yielded significant variation between strains on both antibiotics [vancomycin: F(2, 6) = 24.6, p < 0.002; bacitracin: F(2, 6) = 42.2, p < 0.0003]. A post hoc Tukey test showed that Δ*spt* differed from wild-type and the *spt*-complement strain on both antibiotics significantly at p < 0.05 (n=3 biological replicates, error bars are SD). (B) The indicated strains were labelled with BODIPY-vancomycin and imaged. Scale bar= 5 µm. Representative images from two biological replicates. (C) The indicated strains were grown overnight with or without 0.6 mg/ml deoxycholate before plating on PYE plates (n=2 biological replicates). (D) The expression of genes involved in the unfolded-protein response was measured in the indicated strains by qRT-PCR. Expression of each gene was normalized to the expression in wild-type cells (n=3 biological replicates, error bars are SEM, * p < 0.05, two-tailed T-test).

Given the defects in outer membrane permeability, we hypothesized that sphingolipid depletion may also affect OM protein insertion. In *E. coli*, inhibition of the BAM complex induces the unfolded-protein stress response which includes the induction of DegP chaperone/proteases and proteins involved in OM homeostasis (LolA, BamB, LpxC) (22). Similarly, we observed induction of many of these genes, particularly the lipoprotein transporter *lolA*, in *C. crescentus* upon *spt* deletion (Figure 1D), further supporting a role for sphingolipids in overall OM homeostasis. By contrast, we did not observe any change in the expression of general protein folding chaperones *dsbA* or *fkbP*, which mediate disulfide bond formation and peptidyl-prolyl cis-trans isomerization, respectively (Figure 1D).

*Bioinformatic identification of candidate sphingolipid transport genes in* C. crescentus

The operons responsible for sphingolipid biosynthesis in *C. crescentus* span *ccna_01212* to *ccna_01226*. Within this locus, we identified three genes with homology to well-characterized inner-membrane ABC transporter of the lipopolysaccharide (LPS) transport pathway. The ABC transporter consists of the transmembrane permease subunits LptFG and two cytoplasmic LptB ATPase proteins (Figure 2A). LptC is a periplasmic protein that is anchored in the IM, interacts with LptFG, and passes the extracted LPS to LptA which then delivers LPS to LptDE for insertion into the outer leaflet of the OM (Figure 2A). The sphingolipid locus encodes apparent duplicates of LptF, LptG, and LptC: CCNA_01214, CCNA_01213, and CCNA_01226, respectively (Supplementary Figure 1, Figure 2B). In addition to sequence homology, the Alphafold structures of all three proteins have a high degree of alignment (Figure 2C). For ease of nomenclature, we will refer to these genes as *lptF2*, *lptG2*, and *lptC2* for the remainder of this manuscript.

**Figure 2:**
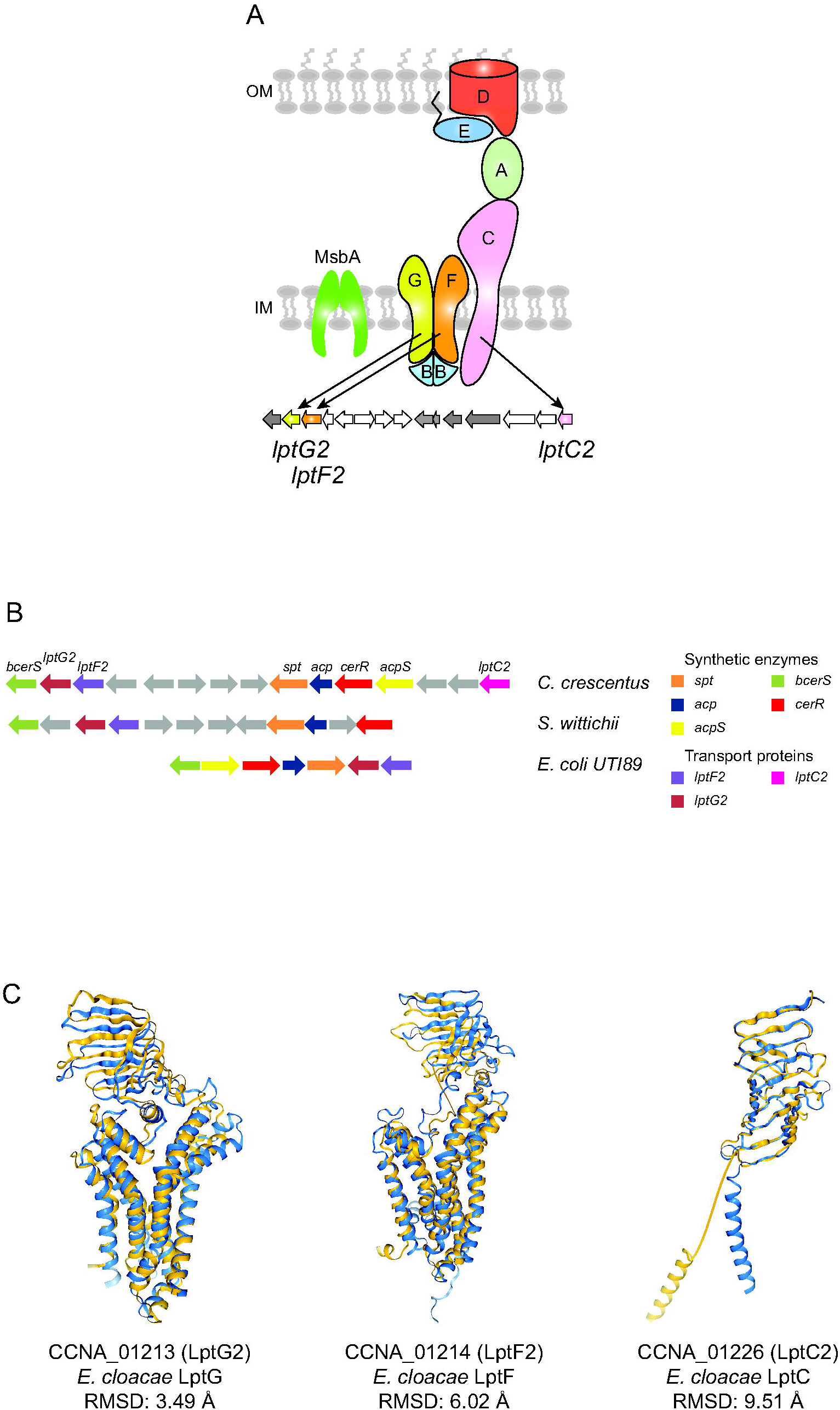
Homologues of LptFGC are present in the sphingolipid locus. (A) This cartoon diagrams the LPS transport pathway. Homologues of LptFGC are encoded within the sphingolipid synthesis locus. (B) The sphingolipid operons from *C. crescentus*, *S. wittichii*, and *E. coli* strain UTI89 are presented. Homologous genes are similarly colored in the three organisms. (C) Foldseek (32) was used to generate an alignment of the Alphafold-predicted structure of the putative sphingolipid transporters (blue) with the corresponding LPS transporters (gold; PDB Accession 6MIT) (25).

Their genomic placement alongside sphingolipid biosynthetic genes suggested that these homologues may have evolved for sphingolipid transport. Supporting this hypothesis, homologues of LptF2/G2 are present in *Sphingomonas* species, which contain sphingolipids instead of LPS. For instance, in *S. wittichii*, the LptFG homologues are found within the same genomic region as the sphingolipid biosynthetic genes, similar to the organization observed in *C. crescentus* (Figure 2B). BLAST analysis further indicates that these homologues represent the sole LptFG proteins encoded in *S. wittichii*, suggesting that they are likely dedicated sphingolipid transporters.

### Genetic evidence for LptF2, LptG2, and LptC2 in lipid transport

Our initial approach to studying the sphingolipid *lpt* genes was to generate individual gene deletions for further characterization. Interestingly, we were unable to generate any of the three deletion mutants, suggesting that these genes were essential (Figures 3A-B). Given that sphingolipids themselves are not required for viability, we hypothesized that deletion of any one of the transporter subunits may lead to toxic accumulation of sphingolipids in the IM. Indeed, we were readily able to generate deletions of *lptF2*, *lptG2*, and *lptC2* in the Δ*spt* background in which sphingolipid synthesis is abolished (Figure 3C). Furthermore, the primary OM sphingolipid in *C. crescentus* is CPG2 (12), which has a high degree of negative charge. We tested whether this anionic headgroup was the source of IM toxicity by attempting to delete these genes in the Δ*cpgB* background which can only synthesize neutral ceramide. As with the Δ*spt* strain, we were able to delete each of the putative transporters in this background (Figure 3D).

**Figure 3:**
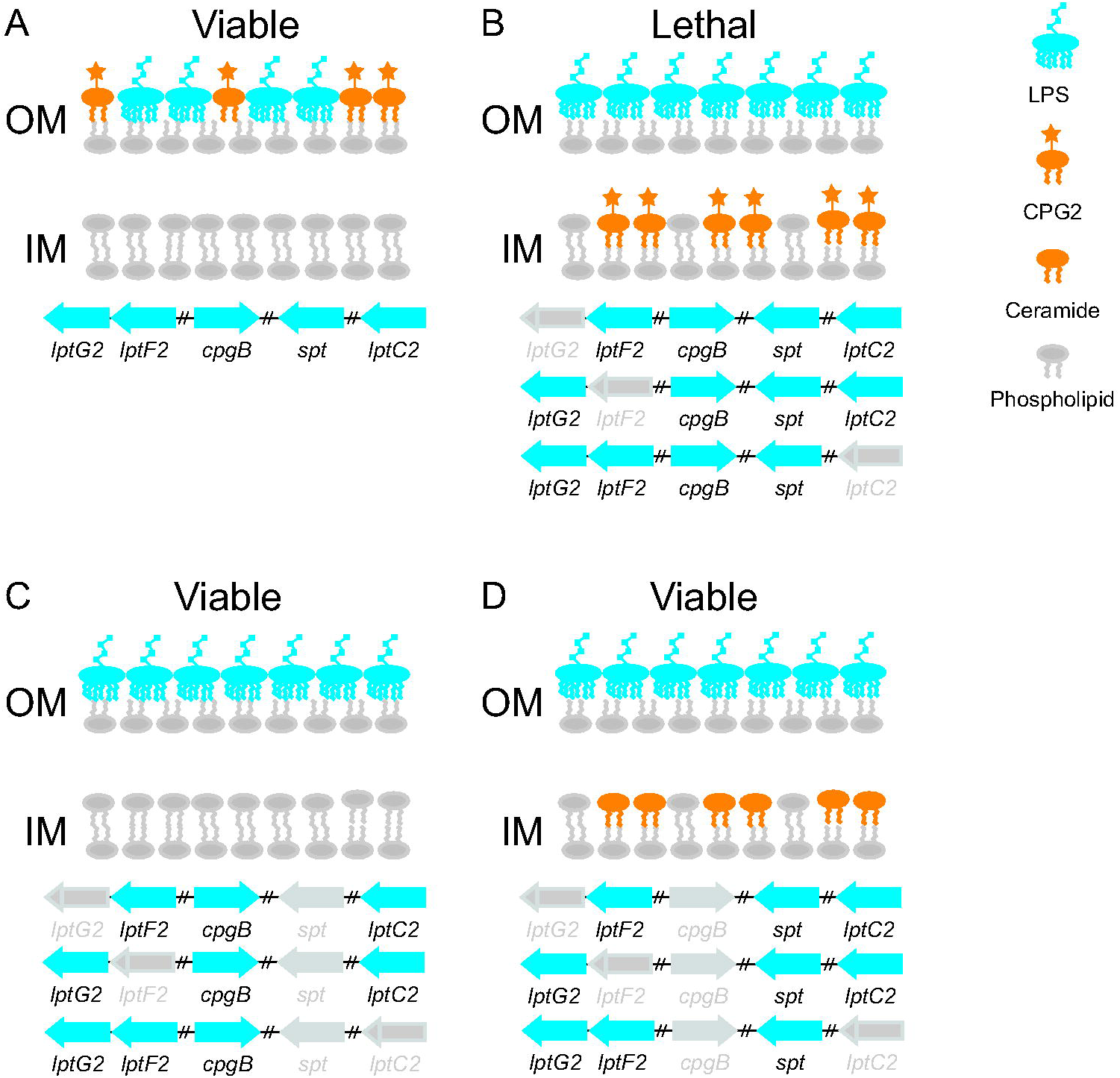
Genetic evidence for LptF2, LptG2 and LptC2 in sphingolipid transport. (A-D) These cartoons illustrate the various attempted deletion mutants, their hypothesized effects on lipid transport, and the viability of the resulting mutant. For each strain, the genes colored in blue are present, whereas the gray genes are deleted. In the presence of both *spt* and *cpgB*, *C. crescentus* produces the anionic sphingolipid CPG2. Deletion of *cpgB* results in the synthesis of neutral ceramide. (B) Under conditions of CPG2 production, we were not able to obtain deletions of *lptF2*, *lptG2*, or *lptC2*. (C-D) Each of these genes was readily deleted in either the *spt* or *cpgB* mutant backgrounds.

### Sphingolipid Lpt proteins form a complex

To determine whether LptF2, LptG2, and LptC2 interact to form a functional complex, we performed a series of pull-down assays. For each transporter subunit, we expressed a C-terminal twin-strep tagged protein while simultaneously expressing the other transporters with a C-terminal FLAG epitope from a chromosomally integrated xylose-inducible locus. Both LptG2 and LptC2 were strep-tagged at the native chromosomal locus. We were unable to generate a native LptF2-strep strain and instead expressed this protein from a chromosomally integrated vanillate-inducible locus. Protein complexes were purified from membranes via the twin-strep tag using Strep-Tactin XT resin, and co-purified proteins were detected by Western blot. LptF2 and LptG2 co-eluted using either protein as the bait (Figures 4A-B). By contrast, we could not detect any interaction between LptC2 and the other transport proteins (Supplemental Figure 2). The genetic data above showed that *lptC2* could only be deleted in a CPG2-deficient background suggesting that this protein is involved in sphingolipid transport. Furthermore, the fact that we were able to generate a natively-tagged LptC2 indicates that the strep-tag does not interfere with the protein’s function. The lack of co-purification between LptC2 and the other transporters may reflect a weaker or transient interaction that was not detected in this assay.

**Figure 4:**
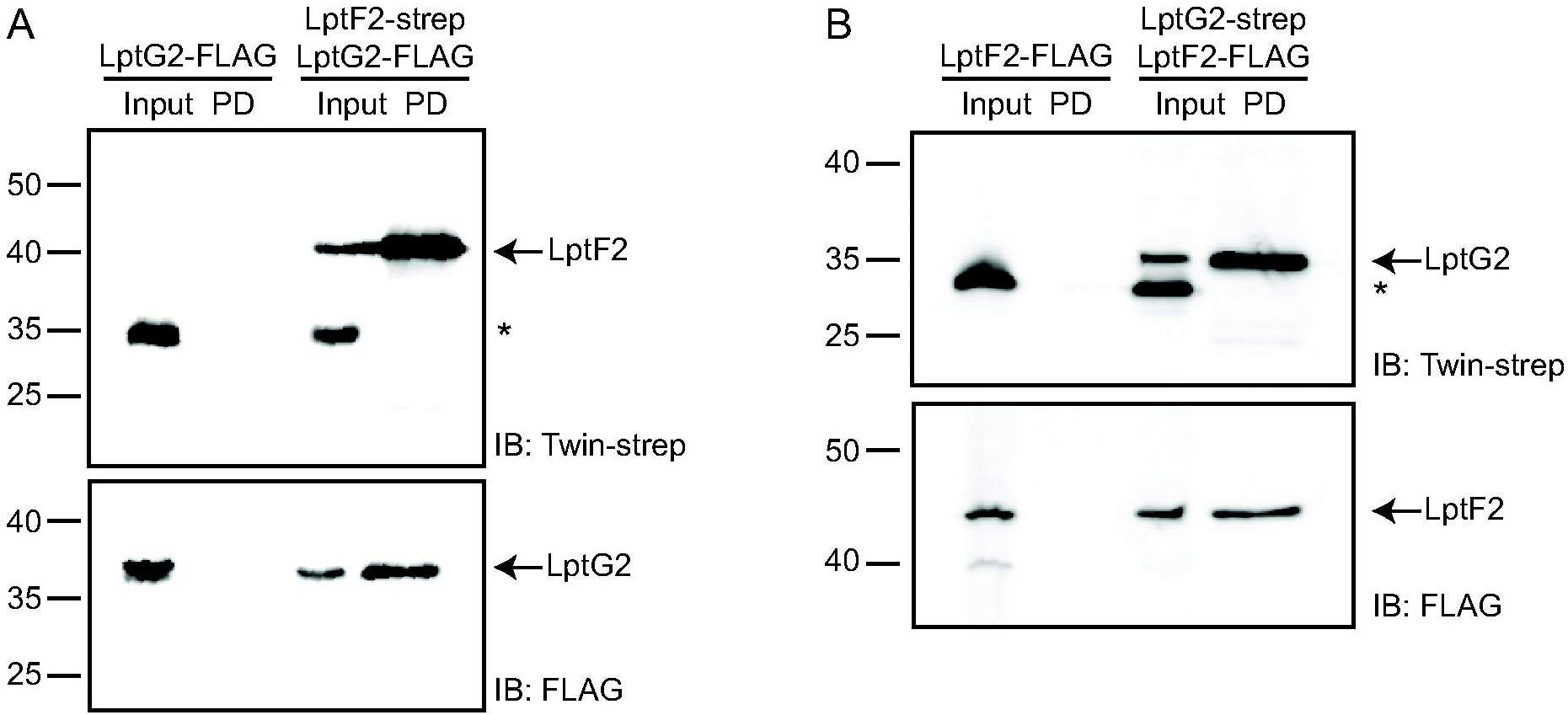
LptF2 and LptG2 protein interactions. LptF2 and LptG2 proteins were expressed with twin-strep tags to enable pull-down experiments. In parallel, the opposite proteins were expressed with FLAG tags to enable detection. As a control, Strep-Tactin XT pull downs were done on strains expressing only the FLAG-tagged protein to ensure there was no non-specific binding. The input lanes are total cell lysates, and the PD (pull-down) lanes show the eluted proteins from the Strep-Tactin XT column. The (*) indicates a non-specific band detected by the anti-strep tag antibody which did not bind to the affinity resin. Both LptF2-strep (A) and LptG2-strep (B) could pull down their respective partners.

### Sphingolipid transport shares a component with LPS transport

The extraction of lipids from the IM for transport to the OM requires energy. For LPS transport, the ATP binding protein LptB binds to the LptFGC complex and drives lipid extraction via ATP hydrolysis (23). In the case of sphingolipid transport, we did not find a duplicate of the *lptB* gene in the sphingolipid locus, or elsewhere on the chromosome. Given the structural similarity between the LPS and sphingolipid transporters, we hypothesized that a single LptB protein might power both inner-membrane complexes. Using epitope-tagged LptB as above, we observed interactions between LptB and both LptF2 and LptG2 by co-purification and western blot (Figures 5A-D).

**Figure 5:**
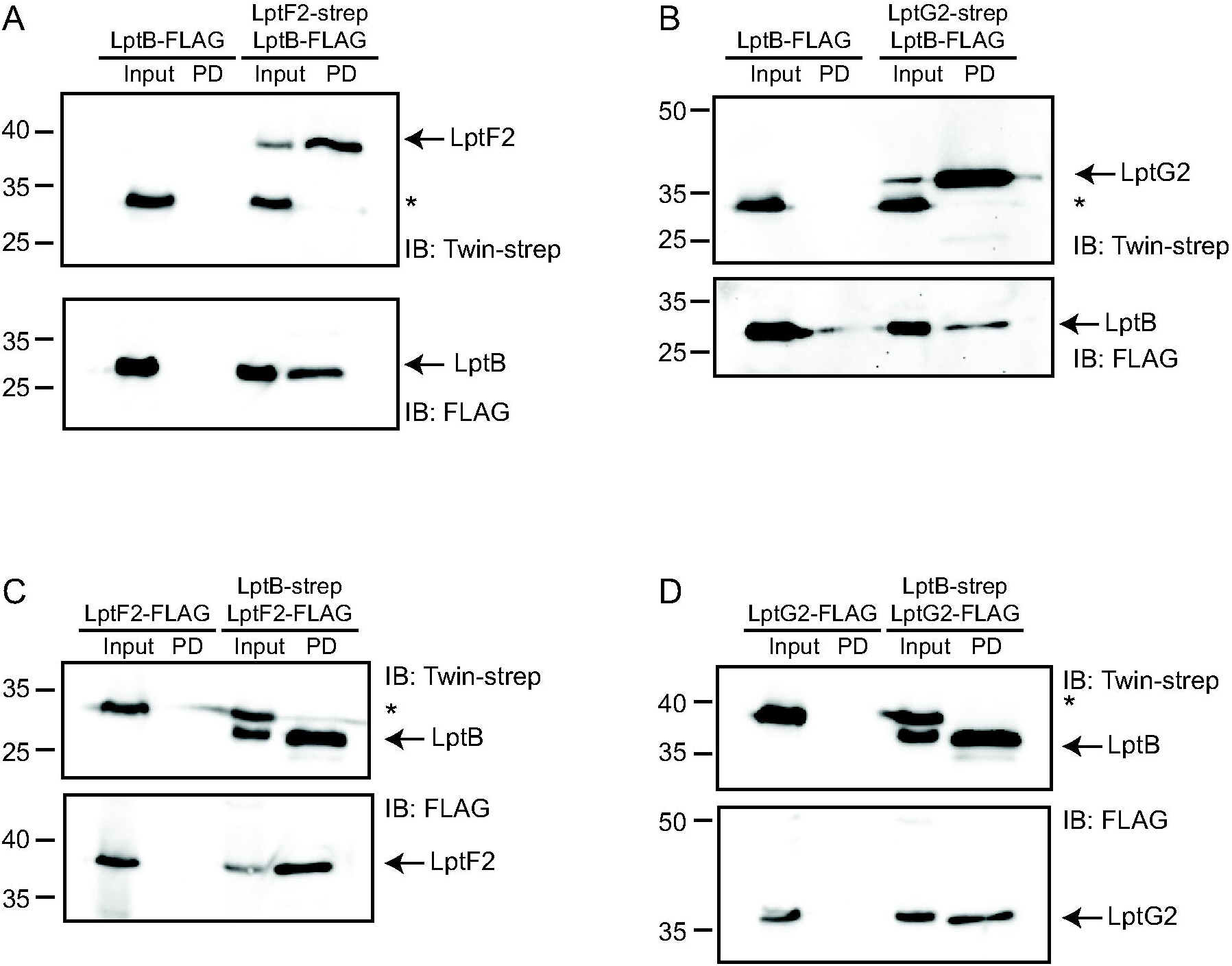
LptF2 and LptG2 form a complex with LptB. Strep-tagged LptF2, LptG2, and LptB were used in pull down assays to detect protein-protein interactions with the corresponding FLAG-tagged proteins. The input lanes are total cell lysates, and the PD (pull-down) lanes show the eluted proteins from the Strep-Tactin XT column. The (*) indicates a non-specific band detected by the anti-strep tag antibody which did not bind to the affinity resin. Both strep-tagged LptF2 (A) and LptG2 (B) were able to pull down LptB. Similarly, strep-tagged LptB was able to pull down LptF2 (C) and LptG2 (D).

Consistent with our biochemical data, the Alphafold 3 (24) predicted structure of the *C. crescentus* LptB_2_F2G2 complex showed a high degree of similarity (RMSD 1.45Å) to the crystal structure of the LptB_2_FG complex from *E. cloacae* (25) (Figure 6A). When we used Alphafold 3 to generate a structure of the complex including LptC2, it likewise aligned well with the *E. cloacae* structure (RMSD 2.18Å) (Figure 6B). We note that the N-terminal tail of LptC, which inserts into the IM and interacts with LptFG, is not well resolved in the crystal structure; we therefore excluded this domain of LptC2 from our model alignment.

**Figure 6:**
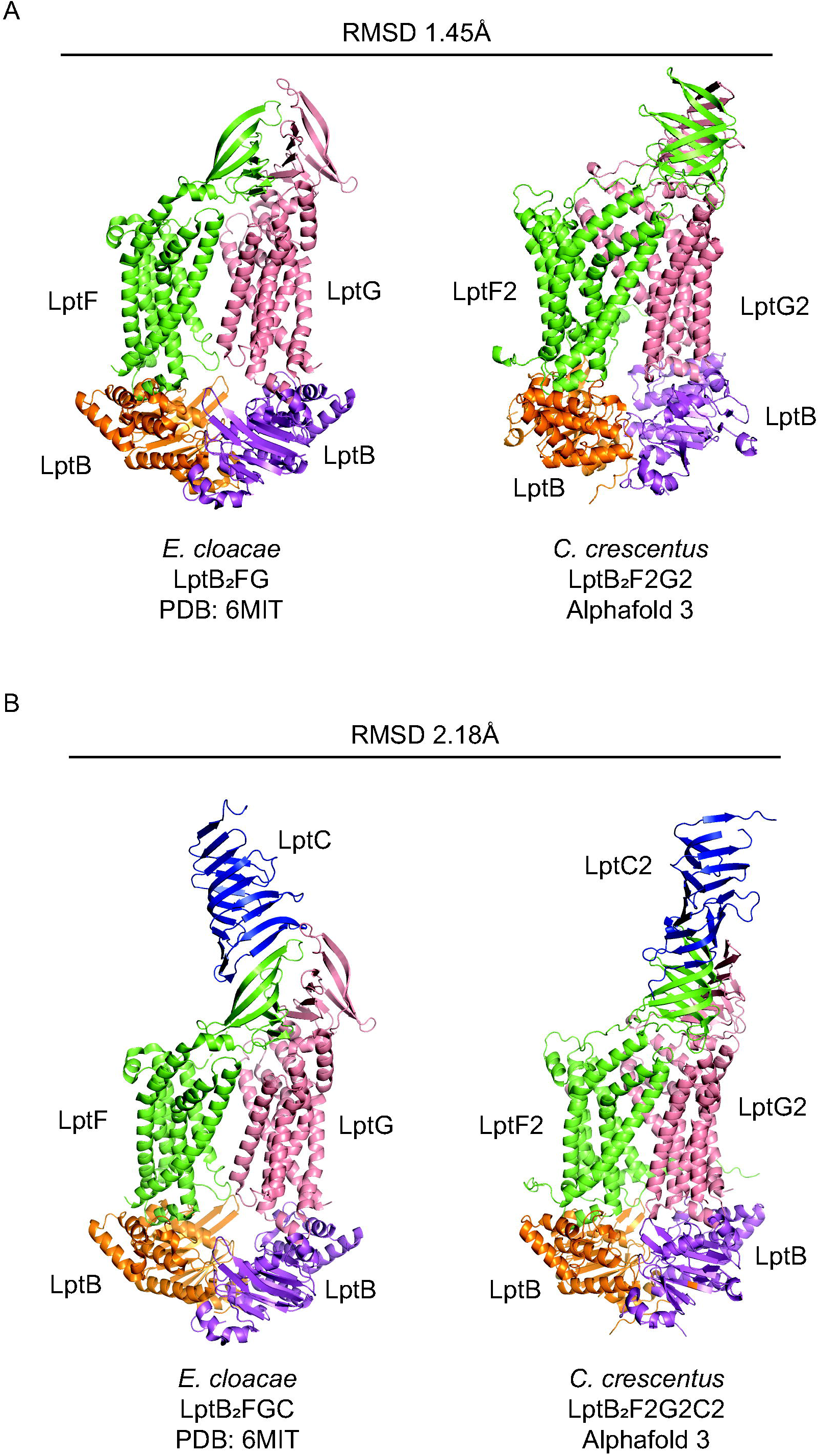
Structural models of the sphingolipid transporter are similar to their LPS counterpart. (A) A model of the *C. crescentus* LptB_2_F2G2 complex was generated with Alphafold 3. The RMSD between this model and the crystal structure of *E. cloacae* LptB_2_FG was determined using the RMSD Calculator (https://www.tamarind.bio). (B) An Alphafold 3 model of *C. crescentus* LptB_2_F2G2C2 was compared to the *E. cloacae* LptB_2_FGC structure. The N-terminal IM tails of LptC/LptC2 were not included in the alignment since these residues were not well resolved in the crystal structure (*E. cloacae* 1-42; *C. crescentus* 1-64).

## Discussion

Sphingolipids (SLs) are increasingly recognized as important components of outer membranes in a wide range of bacterial taxa. While not required for viability, sphingolipid-deficient *C. crescentus* activated OM stress responses and displayed increased sensitivity to antibiotics including bacitracin and vancomycin. Induction of *lolA* in the Δ*spt* strain indicated that loss of sphingolipid production disrupts envelope homeostasis. While we previously demonstrated that *C. crescentus* can tolerate the absence of LPS due to the production of anionic ceramide diphosphoglycerate (CPG2) (12), our findings suggest that even with both LPS and sphingolipids present, these lipids may function together to reinforce the outer membrane against environmental stresses.

Our lab and others have elucidated many of the biochemical pathways required for sphingolipid synthesis and modification, yet the mechanisms by which these lipids are transported across the cell envelope remained poorly understood. In this study, we identified three candidate IM transport proteins encoded within the sphingolipid locus, with close homology to the IM LPS transport proteins, LptFGC. Similar genomic arrangements have been observed in other sphingolipid-producing bacteria, including *S. wittichii* where genes encoding putative LPS transport homologs are positioned adjacent to the *spt* operon.

Genetic evidence suggests that these candidate IM transporters function in the trafficking of sphingolipids. Notably, deletion of the putative transporter genes was only possible in the absence of *spt*. This dependency suggests that the transport machinery functions to transport sphingolipids away from the IM, which would occur downstream of sphingolipid synthesis. Because sphingolipid synthesis starts at the cytoplasmic leaflet of the IM (19), deletion of the transporters resulted in the accumulation of sphingolipids in the IM leading to toxicity. This scenario is similar to LPS biosynthesis, where a defect in transport leads to the accumulation of intermediates in the IM that can disrupt membrane homeostasis (23, 26), prompting the bacterium to downregulate synthesis when transport is impaired. Likewise, in *E. coli*, defects in the OM or in LPS transport trigger compensatory upregulation of LPS biosynthesis genes (27). By analogy, bacterial sphingolipids may be regulated by comparable feedback mechanisms that synchronize their synthesis and translocation across the cell envelope.

Consistent with the genetic data, protein-protein interaction studies demonstrated that CCNA_01213 (LptG2) and CCNA_01214 (LptF2) form a complex. We were not able to observe any interaction between these proteins and CCNA_01226 (LptC2). The fact that *lptC2* could only be deleted in the Δ*spt* background suggests that this protein may be involved in sphingolipid transport. Thus, the lack of co-purification may reflect transient or weak interactions. We do note that while *lptF2* and *lptG2* are present in many sphingolipid loci, particularly in alpha- and gammaproteobacteria (Figure 2B), *lptC2* appears to be restricted to *Caulobacter* species. Therefore, we cannot yet conclude a role for LptC2 in sphingolipid trafficking.

LPS transport is powered by the ATPase LptB, which interacts with LptFG. As there is no homologue of LptB in the sphingolipid locus, we tested and found that LptF2 and LptG2 interact with the sole LptB protein in *C. crescentus*. The association of this complex with LptB supports a model in which LptB/F2/G2 form a functional ABC transporter, with LptF2/G2 as the permease domains and LptB as the ATPase, indicating that sphingolipid trafficking partially co-opts the energy-coupling mechanism of the canonical LPS transport system.

Despite these advances in our understanding of sphingolipid transport, several important questions remain. While our results identify candidate IM transporters and suggest a role for LptB in sphingolipid export, the downstream components responsible for moving sphingolipids across the periplasm and inserting them into the OM remain unknown. In the LPS pathway, the LptA periplasmic bridge and the LptD/E outer membrane complex are essential for lipid translocation (21, 23, 26, 27). We have not found duplications of these genes in the sphingolipid locus of *C. crescentus* or other organisms. One possibility is that, like LptB, sphingolipids share Lpt components with the LPS transport system. Another possibility is that there are yet unidentified proteins required for sphingolipid trafficking. Organisms of the *Sphingomonas* genus do not produce any LPS and lack the key LPS synthesis enzymes including LpxC, LpxB, and WaaA. Nevertheless, these organisms encode homologues of the entire Lpt transport system including LptA, LptE, and LptD, perhaps to traffic sphingolipids. We therefore favor a model where LptF2 and LptG2 utilize this system.

A second important question is whether this transport mechanism is used by all sphingolipid-producing bacteria. Bacteroidetes and Myxococci are well characterized sphingolipid producers that encode only a single copy of *lptFG*. This suggests that either the canonical Lpt pathway can transport both LPS and sphingolipids in these organisms, or there is an alternative transport pathway. For example, *Myxococus xanthus* encodes an MmpL family transporter immediately downstream of *spt*. This family of transporters exports lipids across the cell envelope (28) and its expression is regulated in a similar manner to sphingolipid synthesis across the *Myxococcus* life cycle (29, 30). Therefore, we hypothesize that sphingolipid transport mechanisms will vary between taxa.

## Materials and Methods

### Bacterial strains, plasmids and growth conditions

Strains, plasmids, and primers used in this study are listed in Supplemental Tables S1–S3, and detailed descriptions of strain construction are provided in the Supplementary Information. *Caulobacter crescentus* wild-type strain NA1000 and derivative strains were routinely cultured at 30 °C in peptone yeast extract (PYE) medium, while *Escherichia coli* strains were grown at 37 °C in Luria–Bertani (LB) medium. Antibiotics were added as required at the following concentrations: tetracycline at 12 µg mL^-1^ for *E. coli* in broth and agar, and at 1 µg mL^-1^ (broth) or 2 µg mL^-1^ (agar) for *C. crescentus*; spectinomycin at 50 µg mL^-1^ for *E. coli* in broth and agar, and at 25 µg mL^-1^ (broth) or 100 µg mL^-1^ (agar) for *C. crescentus*. Gene expression in *C. crescentus* was induced by the addition of 0.5 mM vanillate or 0.075% (w/v) xylose.

### Antibiotic sensitivity assays

The indicated bacterial strains were grown overnight in PYE with the appropriate inducers. The cultures were normalized to an OD_660_ of 0.2 and 250 µl of culture was added to 4 ml of PYE with 0.75% (w/v) agar. The mixture was overlayed onto a PYE-agar plate and allowed to solidify and dry. Sterile Whatman filter paper discs (6 mm diameter) were treated with 20 µl of antibiotics (vancomycin 20 mg ml^-1^; bacitracin 10 mg ml^-1^) or a water control. After drying, the filter discs were laid on top of the agar plates and incubated overnight at 30 °C before measuring the zone of clearance using ImageJ (31).

### BODIPY-vancomycin labeling

Bacterial strains were grown overnight in PYE with the relevant inducers. The cultures were diluted 1:1 with fresh PYE. The vancomycin staining solution was prepared by mixing equal volumes of 1.5 mg ml^-1^ vancomycin with 1.5 mg ml^-1^ BODIPY-vancomycin (Thermo Scientific). 2 µl of the staining solution was added to 1 ml of the diluted bacteria and the cells were incubated with the dye for 10 minutes at room temperature in the dark. The cells were washed three times in PBS before imaging in phase contrast and fluorescence (GFP filter set). Imaging was performed on a Nikon Ti-E inverted microscope equipped with a Prior Lumen 220PRO illumination system, CFI Plan Apochromat 100X oil immersion objective (NA 1.45, WD 0.13 mm), Nikon Qi2 camera, and NIS Elements v, 4.20.01 for image acquisition.

### Deoxycholate sensitivity assay

The indicated bacterial strains were grown overnight in PYE with the appropriate inducers, with or without 0.6 mg ml^-1^ sodium deoxycholate. Following overnight growth, serial dilutions of each culture were made in plain PYE and 5 µl samples were spotted onto PYE plates to monitor colony formation.

### Quantitative reverse transcriptase PCR (qRT-PCR)

RNA was extracted from bacterial cultures using the Qiagen RNeasy kit. Following DNase digestion, RNA (25 ng μl^-1^) was reverse transcribed using the high-capacity cDNA reverse transcription kit (Applied Biosystems). A total of 0.5 μl of cDNA was used as a template in a 10-μl qPCR performed with iTaq Universal SYBR Green Supermix (Bio-Rad). qPCR was performed on an ABI QuantStudio 6 instrument using the ΔΔCT (where CT is threshold cycle) method. *rpoD* expression was used as the loading control. qPCR primer sequences are available in Supplemental Table S3.

### Protein structure prediction and analysis

The proteins sequences of LptF2/G2/C2/B were uploaded to the Alphafold 3 server (https://alphafoldserver.com/) to generate models of the transporter complex. The top-scoring models were compared to the crystal structure of the *E. cloacae* LptB_2_FGC structure (PDB: 6MIT (25)) using the RMSD Calculator (https://www.tamarind.bio/).

### Pull down assay

Strains expressing twin strep-tagged proteins were grown in PYE medium at 30 °C, back-diluted 1:2000 into 1 L cultures and allowed to express overnight. Cells were harvested by centrifugation at 10,000 × g for 10 min at 4 °C and resuspended in 25 ml of membrane lysis buffer (1x PBS, 10% glycerol) supplemented with lysozyme (1 mg mL^-1^). Cells were lysed by French press (three passes at 10,000 psi), and unbroken cells were removed by centrifugation at 15,000 × g for 10 min. Total membranes were collected by ultracentrifugation at 150,000 × g for 1 hour at 4 °C and solubilized in 10 ml of membrane resuspension buffer (50 mM Tris pH 8.0, 500 mM NaCl, 10% glycerol, 25 mM n-dodecyl-β-D-maltoside (DDM)) for 2 hours at 4 °C. Solubilized membranes were subjected to a second ultracentrifugation step (150,000 × g for 1 h), after which the supernatant was collected and the pellet discarded.

Strep-Tactin XT 4Flow columns (1 ml bed volume, IBA Lifesciences) were equilibrated in 2 ml membrane resuspension buffer (50 mM Tris pH 8.0, 500 mM NaCl, 10% glycerol) containing 25 mM DDM, and the solubilized protein fraction was poured into the column. Flow-through fractions were collected, and the resin was washed with eight column volumes of wash buffer (50 mM Tris pH 8.0, 500 mM NaCl, 10% glycerol) containing 0.5 mM DDM. Bound proteins were eluted using biotin-containing elution buffer (50 mM Tris pH 8.0, 500 mM NaCl, 10% glycerol, 50 mM biotin) supplemented with 0.5 mM DDM. Total lysate, flow-through, wash, and elution fractions were analyzed by SDS-PAGE without boiling (except total lysate).

### Western blotting

Proteins were separated by SDS-PAGE on 12% polyacrylamide gels and transferred to nitrocellulose membranes. Membranes were blocked with 5% non-fat milk in wash buffer (20 mM Tris-HCl, pH 7.5, 50 mM NaCl, 2.5 mM EDTA, and 0.1% Tween-20) and probed with primary antibodies against the Strep tag (Abcam, anti–Strep-tag II; ab183907; 1:5,000) or FLAG tag (Proteintech; 20543-1-AP; 1:1,000). After washing (3X, 10 min per wash), membranes were incubated with horseradish peroxidase–conjugated secondary antibodies (1:5,000), washed again (3X, 10 minutes per wash), and proteins were detected with Super signal West Pico Plus chemiluminescent substrate (Thermo Fisher Scientific). Signals were visualized using a Bio-Rad ChemiDoc MP imaging system.

## Data availability

All of the data for this work is contained within the manuscript.

## Acknowledgements

We thank Suzanne Letham (University of Oxford) for assistance in developing the co-purification methods and Marcin Grabowicz (Emory University) for helpful discussions.

## Funding

Funding was provided by National Science Foundation grant MCB-2224195 (E.A.K.).

## Conflict of interest

The authors declare that they have no conflicts of interest with the contents of this article.

## Author CrediT statement

UGC, GLI, and EAK conceptualization; CGU, GLI, and EAK methodology; CGU, GLI, and EAK investigation; CGU and EAK writing-original draft; CGU and EAK visualization; GLI writing-review & editing; EAK supervision.

## Notes

### Competing Interest Statement

The authors have declared no competing interest.

## References

1. Ruiz N, Wu T, Kahne D, Silhavy TJ. 2006. Probing the barrier function of the outer membrane with chemical conditionality. ACS Chem Biol 1:385–95.

2. Nikaido H. 2003. Molecular basis of bacterial outer membrane permeability revisited. Microbiol Mol Biol Rev 67:593–656.

3. Carey AB, Ashenden A, Koper I. 2022. Model architectures for bacterial membranes. Biophys Rev 14:111–143.

4. Raetz CR, Whitfield C. 2002. Lipopolysaccharide endotoxins. Annu Rev Biochem 71:635–700.

5. Birtles D, Fenn KL, Machin JM, Radford SE, Ranson NA. 2026. Integration of membrane proteins into the outer membrane of diderm bacteria by the BAM complex. Chem Rev doi:10.1021/acs.chemrev.5c00764.

6. Nikaido H. 1989. Outer membrane barrier as a mechanism of antimicrobial resistance. Antimicrob Agents Chemother 33:1831–6.

7. Schroder NW, Eckert J, Stubs G, Schumann RR. 2008. Immune responses induced by spirochetal outer membrane lipoproteins and glycolipids. Immunobiology 213:329–40.

8. Kawasaki S, Moriguchi R, Sekiya K, Nakai T, Ono E, Kume K, Kawahara K. 1994. The cell envelope structure of the lipopolysaccharide-lacking gram-negative bacterium *Sphingomonas paucimobilis*. J Bacteriol 176:284–90.

9. Moffatt JH, Harper M, Harrison P, Hale JD, Vinogradov E, Seemann T, Henry R, Crane B, St Michael F, Cox AD, Adler B, Nation RL, Li J, Boyce JD. 2010. Colistin resistance in *Acinetobacter baumannii* is mediated by complete loss of lipopolysaccharide production. Antimicrob Agents Chemother 54:4971–7.

10. Peng D, Hong W, Choudhury BP, Carlson RW, Gu XX. 2005. *Moraxella catarrhalis* bacterium without endotoxin, a potential vaccine candidate. Infect Immun 73:7569–77.

11. Steeghs L, de Cock H, Evers E, Zomer B, Tommassen J, van der Ley P. 2001. Outer membrane composition of a lipopolysaccharide-deficient *Neisseria meningitidis* mutant. EMBO J 20:6937–45.

12. Zik JJ, Yoon SH, Guan Z, Stankeviciute Skidmore G, Gudoor RR, Davies KM, Deutschbauer AM, Goodlett DR, Klein EA, Ryan KR. 2022. *Caulobacter* lipid A is conditionally dispensable in the absence of fur and in the presence of anionic sphingolipids. Cell Rep 39:110888.

13. Stankeviciute G, Guan Z, Goldfine H, Klein EA. 2019. *Caulobacter crescentus* adapts to phosphate starvation by synthesizing anionic glycoglycerolipids and a novel glycosphingolipid. mBio 10:e00107–19.

14. Olea-Ozuna RJ, Poggio S, Bergstrom E, Osorio A, Elufisan TO, Padilla-Gomez J, Martinez-Aguilar L, Lopez-Lara IM, Thomas-Oates J, Geiger O. 2024. Genes required for phosphosphingolipid formation in *Caulobacter crescentus* contribute to bacterial virulence. PLoS Pathog 20:e1012401.

15. Olea-Ozuna RJ, Poggio S, EdBergstrom, Quiroz-Rocha E, Garcia-Soriano DA, Sahonero-Canavesi DX, Padilla-Gomez J, Martinez-Aguilar L, Lopez-Lara IM, Thomas-Oates J, Geiger O. 2021. Five structural genes required for ceramide synthesis in *Caulobacter* and for bacterial survival. Environ Microbiol 23:143–159.

16. Dhakephalkar T, Guan Z, Klein EA. 2025. CpgD is a phosphoglycerate cytidylyltransferase required for ceramide diphosphoglycerate synthesis. J Biol Chem 301:110386.

17. Dhakephalkar T, Stukey GJ, Guan Z, Carman GM, Klein EA. 2023. Characterization of an evolutionarily distinct bacterial ceramide kinase from *Caulobacter crescentus*. J Biol Chem 299:104894.

18. Stankeviciute G, Tang P, Ashley B, Chamberlain JD, Hansen MEB, Coleman A, D’Emilia R, Fu L, Mohan EC, Nguyen H, Guan Z, Campopiano DJ, Klein EA. 2022. Convergent evolution of bacterial ceramide synthesis. Nat Chem Biol 18:305–312.

19. Uchendu CG, Guan Z, Klein EA. 2024. Spatial organization of bacterial sphingolipid synthesis enzymes. J Biol Chem 300:107276.

20. Doerrler WT, Raetz CR. 2002. ATPase activity of the MsbA lipid flippase of *Escherichia coli*. J Biol Chem 277:36697–705.

21. Botos I, Majdalani N, Mayclin SJ, McCarthy JG, Lundquist K, Wojtowicz D, Barnard TJ, Gumbart JC, Buchanan SK. 2016. Structural and functional characterization of the LPS transporter LptDE from Gram-negative pathogens. Structure 24:965–976.

22. Hews CL, Cho T, Rowley G, Raivio TL. 2019. Maintaining integrity under stress: envelope stress response regulation of pathogenesis in Gram-negative bacteria. Front Cell Infect Microbiol 9:313.

23. Ruiz N, Gronenberg LS, Kahne D, Silhavy TJ. 2008. Identification of two inner-membrane proteins required for the transport of lipopolysaccharide to the outer membrane of *Escherichia coli*. Proc Natl Acad Sci U S A 105:5537–42.

24. Abramson J, Adler J, Dunger J, Evans R, Green T, Pritzel A, Ronneberger O, Willmore L, Ballard AJ, Bambrick J, Bodenstein SW, Evans DA, Hung CC, O’Neill M, Reiman D, Tunyasuvunakool K, Wu Z, Zemgulyte A, Arvaniti E, Beattie C, Bertolli O, Bridgland A, Cherepanov A, Congreve M, Cowen-Rivers AI, Cowie A, Figurnov M, Fuchs FB, Gladman H, Jain R, Khan YA, Low CMR, Perlin K, Potapenko A, Savy P, Singh S, Stecula A, Thillaisundaram A, Tong C, Yakneen S, Zhong ED, Zielinski M, Zidek A, Bapst V, Kohli P, Jaderberg M, Hassabis D, Jumper JM. 2024. Accurate structure prediction of biomolecular interactions with AlphaFold 3. Nature 630:493–500.

25. Owens TW, Taylor RJ, Pahil KS, Bertani BR, Ruiz N, Kruse AC, Kahne D. 2019. Structural basis of unidirectional export of lipopolysaccharide to the cell surface. Nature 567:550–553.

26. Sperandeo P, Lau FK, Carpentieri A, De Castro C, Molinaro A, Deho G, Silhavy TJ, Polissi A. 2008. Functional analysis of the protein machinery required for transport of lipopolysaccharide to the outer membrane of *Escherichia coli*. J Bacteriol 190:4460–9.

27. Sperandeo P, Cescutti R, Villa R, Di Benedetto C, Candia D, Deho G, Polissi A. 2007. Characterization of *lptA* and *lptB*, two essential genes implicated in lipopolysaccharide transport to the outer membrane of *Escherichia coli*. J Bacteriol 189:244–53.

28. Viljoen A, Dubois V, Girard-Misguich F, Blaise M, Herrmann JL, Kremer L. 2017. The diverse family of MmpL transporters in mycobacteria: from regulation to antimicrobial developments. Mol Microbiol 104:889–904.

29. Ahrendt T, Wolff H, Bode HB. 2015. Neutral and phospholipids of the *Myxococcus xanthus* lipodome during fruiting body formation and germination. Appl Environ Microbiol 81:6538–47.

30. Muller FD, Treuner-Lange A, Heider J, Huntley SM, Higgs PI. 2010. Global transcriptome analysis of spore formation in *Myxococcus xanthus* reveals a locus necessary for cell differentiation. BMC Genomics 11:264.

31. Schindelin J, Arganda-Carreras I, Frise E, Kaynig V, Longair M, Pietzsch T, Preibisch S, Rueden C, Saalfeld S, Schmid B, Tinevez JY, White DJ, Hartenstein V, Eliceiri K, Tomancak P, Cardona A. 2012. Fiji: an open-source platform for biological-image analysis. Nat Methods 9:676–82.

32. van Kempen M, Kim SS, Tumescheit C, Mirdita M, Lee J, Gilchrist CLM, Soding J, Steinegger M. 2024. Fast and accurate protein structure search with Foldseek. Nat Biotechnol 42:243–246.

