## Supplemental methods for "Homologues of the inner-membrane LPS transport proteins are required for sphingolipid transport in *Caulobacter crescentus*"

**Supplementary Methods**

*Cloning methods*

Creation of the various *C. crescentus* deletion strains was done by double-homologous recombination using both positive and negative selection. The suicide plasmid pNPTS138 has both a kanamycin resistance cassette as well as the *sacB* gene which is toxic in the presence of sucrose. Genomic fragments (~500 bp) upstream and downstream of the targeted gene were ligated in tandem in the pNPTS138 vector. The plasmid was transformed into *C. crescentus* and recombinants were selected on kanamycin plates. Individual colonies were grown overnight in PYE (without antibiotic) and streaked out onto PYE-3% sucrose plates to recover colonies that performed the second recombination. Colonies were screened for the gene deletion by PCR and streaked out onto plain PYE and PYE-kanamycin plates to confirm the loss of the plasmid backbone.

Deletion of *ccna_01213* (*lptG2*) done by PCR amplifying the upstream (EK1207/1208) and downstream (EK1209/1210) homology fragments from NA1000 genomic DNA. The fragments were stitched together by overlap PCR. The final purified PCR product was cut with HindIII and EcoRI and ligated into the HindIII/EcoRI site of pNPTS138. The assembled plasmid was electroporated into the desired strain followed by selection on PYE-kanamycin plates. An individual colony was grown overnight in PYE and streaked onto PYE-3% sucrose plates. Colonies were screened for the *ccna_01213* deletion with primers EK S275/S276 (wild-type 2 kb; deletion 1.2 kb).

Deletion of *ccna_01214* (*lptF2*) done by PCR amplifying the upstream (EK1507/1508) and downstream (EK1509/1510) homology fragments from NA1000 genomic DNA. The fragments were stitched together by overlap PCR. The final purified PCR product was cut with HindIII and EcoRI and ligated into the HindIII/EcoRI site of pNPTS138. The assembled plasmid was electroporated into the desired strain followed by selection on PYE-kanamycin plates. An individual colony was grown overnight in PYE and streaked onto PYE-3% sucrose plates. Colonies were screened for the *ccna_01214* deletion with primers EK S313/S314 (wild-type 2 kb; deletion 1.2 kb).

Deletion of *ccna_01226* (*lptC2*) done by PCR amplifying the upstream (EK1583/1584) and downstream (EK1585/1586) homology fragments from NA1000 genomic DNA. The fragments were stitched together by overlap PCR. The final purified PCR product was cut with HindIII and EcoRI and ligated into the HindIII/EcoRI site of pNPTS138. The assembled plasmid was electroporated into the desired strain followed by selection on PYE-kanamycin plates. An individual colony was grown overnight in PYE and streaked onto PYE-3% sucrose plates. Colonies were screened for the *ccna_01226* deletion with primers EK S327/S328 (wild-type 1.4 kb; deletion 1.2 kb).

All of the plasmids for generating the twin-strep-tagged alleles of *lptF2*, *lptG2*, *lptC2*, and *lptB* were synthesized by Genscript (Piscataway, NJ). The tag used in all constructs encoded the peptide: SAWSHPQFEKGGGSGGGSGGSAWSHPQFEK. For native tagging of *lptG2* (*ccna_01213*) and *lptC2* (*ccna_01226*), 500 bp upstream and downstream of the respective stop codons were cloned into plasmid pNPTS138. The twin-strep tag was added at the C-terminus. The recombinant plasmid was introduced into Caulobacter crescentus, and allelic exchange was performed using the sacB-based counterselection system to generate the desired strain. Strep-tagged alleles were identified by PCR: *lptG2* primers EK1209/EK1449 (WT 260 bp, strep-tagged 350 bp) and *lptC2* primers EK1585/1586 (WT 520 bp, strep-tagged 610 bp).

Vanillate-inducible plasmids for strep-tagged LptF2 and LptB were generated by synthesizing a C-terminal twin-strep tag of the respective gene and inserting the coding sequence into the NdeI/NheI site of pVCFPC-1. Plasmids were transformed into *C. crescentus* and integration into the *vanA* locus was confirmed by PCR with primers EK S11/S67 (1.7 kb product).

Xylose-inducible plasmids for FLAG-tagged LptF2, LptG2, LptC2, and LptB were cloned by PCR amplifying the respective genes and inserting the product into the NdeI/NheI site of pXCHYC-5. The following primer pairs were used for cloning: *lptG2* (EK 1211/1212), *lptF2* (EK 1593/1594), *lptC2* (EK 1597/1598), *lptB* (EK 1907/1908).

**Supplemental Table S1: Strains used in this study**

| **Strain** | **Genotype** | **Construction** | **Source** |
| --- | --- | --- | --- |
| *C. crescentus* | | | |
| NA1000 | Synchronizeable variant of wild-type *C. crescentus* strain CB15 |  | (1) |
| CU113 | *ccna_01213 twin strep (C-terminal)* | Transformation of NA1000 with pCU111 | This Study |
| CU114 | *ccna_01226 twin strep (C-terminal)* | Transformation of NA1000 with pCU112 | This Study |
| CU141 | *ccna_01213-twin strep; PxylX::ccna_01214-FLAG* | Transformation of CU113 with pCU95 | This study |
| CU142 | *ccna_01213-twin strep; PxylX::ccna_01226-FLAG* | Transformation of CU113 with pCU93 | This study |
| CU143 | *ccna_01226-twin strep; PxylX::ccna_01214-FLAG* | Transformation of CU114 with pCU95 | This study |
| CU156 | *ccna_01213-twin strep; Pxyl:: LptB-FLAG* | Transformation of CU113 with pCU155 | This study |
| CU162 | *PxylX::LptB-FLAG;*  *PvanA::ccna_01214-twin strep* | Transformation of CU158 with pCU160 | This study |
| CU163 | *PxylX::ccna_01213-FLAG;*  *PvanA::ccna_01214-twin strep* | Transformation of CU90 with pCU160 | This study |
| CU164 | *PxylX::ccna_01213-FLAG;*  *PvanA::LptB-twin strep* | Transformation of CU90 with pCU161 | This study |
| CU165 | *PxylX::ccna_01214-FLAG;*  *PvanA::LptB-twin strep* | Transformation of CU115 with pCU161 | This Study |
| GS32 | *Δccna_01220 (spt)* |  | (2) |
| CU19 | Δ*ccna_01220*;  P*vanA*:: *ccna_01220-FLAG* | Transformation of GS32 with pCU16 | (3) |
| CU127 | *Δccna_01218* |  | (4) |
| CU90 | *PxylX::ccna_01213-FLAG* | Transformation of NA1000 with pCU88 | This study |
| CU115 | *PxylX::ccna_01214-FLAG* | Transformation of NA1000 with pCU95 | This study |
| CU158 | *PxylX::LptB-FLAG* | Transformation of NA1000 with pCU155 | This study |
| *E. coli* | | | |
| DH5a | Cloning strain |  | Thermo Scientific |
| S17 | λ−pir cloning strain, Spec^R^ |  | (5) |

**Supplemental Table S2: Plasmids used in this study**

| **Name** | **Description** | **Source** |
| --- | --- | --- |
| pVCFPC-1 | Vanillate-inducible expression, Spect^R^ | (6) |
| pXCHYC-5 | Xylose-inducible expression, Tet^R^ | (6) |
| pNPTS138 | *sacB*-containing suicide vector used for double homologous recombination, Kan^R^ | Alley, M.R.K. (unpublished) |
| pCU2 | pNPTS based plasmid for deleting *ccna 01213* | This study |
| pCU9 | pNPTS based plasmid for deleting *ccna 01214* | This study |
| pCU35 | pNPTS based plasmid for deleting *ccna 01226* | This study |
| pCU111 | pNPTS based plasmid for generating *ccna_01213-twin strep,* Kan^R^ | This study |
| pCU112 | pNPTS based plasmid for generating *ccna_01226-twin strep,* Kan^R^ | This study |
| pCU88 | pXCHYC-5 based plasmid for *ccna_01213-Flag* expression, Tet^R^ | This study |
| pCU93 | pXCHYC-5 based plasmid for *ccna_01226-Flag* expression, Tet^R^ | This study |
| pCU95 | pXCHYC-5 based plasmid for *ccna_01214-Flag* expression, Tet^R^ | This study |
| pCU155 | pXCHYC-5 based plasmid for *LptB-Flag* expression, Tet^R^ | This study |
| pCU160 | pVCFPC-1-based plasmid for *ccna_01214-twin strep* expression, Spect^R^ | This study |
| PCU161 | pVCFPC-1-based plasmid for L*ptB-twin strep* expression, Spect^R^ | This Study |

**Supplemental Table S3: Primers used in this study**

| **Name** | **Sequence** |
| --- | --- |
| EK1207 | tactaagcttACCCTGCTCAGCGAGTTTTA |
| EK1208 | cgtcaatgcGCCTCGGTCCAGGATGTC |
| EK1209 | gaccgaggcGCATTGACGCCCTTCCTG |
| EK1210 | tactgaattcTCGTTGATCGACAGGTTGAA |
| EK1211 | tactcatATGAAGCTGCAGCTCTATGTCCTG |
| EK1212 | tactgctagcTTActtgtcatcgtcatccttgtagtcGCCCTCCAGGGCCAGGAA |
| EK1449 | tactgtcgacGAAGAACGGATTGGTCTTGG |
| EK1507 | tactgaattcCAAGACCCAGGTCGACTATGA |
| EK1508 | tccagatcgTCACGATGACGATGAACAGG |
| EK1509 | tcatcgtgaCGATCTGGATCCCCTTCAC |
| EK1510 | tactaagcttATGCCCACGATCCGGTAG |
| EK1583 | tactgaattcCGGTTCTGGGCGTAGAGG |
| EK1584 | acctcatcgCCTTTAGAACGCCGGTGAAG |
| EK1585 | ttctaaaggCGATGAGGTGAAGATCACCAG |
| EK1586 | tactaagcttGTGATGACCGCGTCGTAGC |
| EK1593 | tactcatATGAGCGCCAAGGACCTGC |
| EK1594 | tactgctagcTTActtgtcatcgtcatccttgtagtcTGCGGCTTGTCGCTTCTTCG |
| EK1597 | tactcatATGCTCACCCAGTCCGATGG |
| EK1598 | tactgctagcTTActtgtcatcgtcatccttgtagtcGGGTCGCGGCTCCAGCCG |
| EK1907 | tactcatATGACCCTGACCTCCAAAGACATG |
| EK1908 | tactgctagcTTActtgtcatcgtcatccttgtagtcGTCGCCCAGGCGCGGCTC |
| EK S11 | cagccttggccacggtttcggtacc |
| EK S67 | atgccgtttgtgatggcttccatgtcg |
| EK S275 | GATCCGGTCGTCTCCAAAG |
| EK S276 | TTGTAGGCGAAGAGGTCCTG |
| EK S313 | GCGTTCTCGGACGTGAGT |
| EK S314 | TGGCGCCGATGACTAGGT |
| EK S327 | CATCGACATGACGAAATCCA |
| EK S328 | TGACCAGATCGAACAGCTTG |

*qRT-PCR primers*

| **Gene target** | **Forward primer** | **Reverse primer** |
| --- | --- | --- |
| *rpoD* (*ccna_03142*) | CTCTATGCGATCAACAAGCG | ATAGGCCTTGAGGAACTCGC |
| *bamB* (*ccna_01725*) | ATGAACTCCAAGCGTTCCGT | TCTTGTCGAAGGGGTTGAGC |
| *degP1* (*ccna_02846*) | ATCTGAACGACACCACCTCG | GATCACCCGGCCATAGATGT |
| *degP2* (*ccna_01341*) | CTGGTCTGAAGGAAGGCGAC | CCGACCTTGTGGGTTCCAAT |
| *degP3* (*ccna_00150*) | TCAGGGATCCGAGACCGAC | TGGGAATACAAGCTCGACGAC |
| *dsbA* (*ccna_00378*) | GGTCGTGAGGACCTCGAAAT | GCCTAGTCTTCCGCGAGTTT |
| *fkbP* (*ccna_02889*) | CAGCATCCCCTCCAGCTTC | ACGAGTGGACGCTCTTCATC |
| *lolA* (*ccna_03820*) | GGACGGTCACCGACATCAAG | ACGAGGATCGGTCAGGACAA |
| *lpxC* (*ccna_02064*) | TTGTTGATCACCGTGCCGAG | CCGACATGGGGATCGTCTTT |

**Supplemental Figure Legends**

**Supplemental Figure 1: Sequence homology between Lpt homologues.** The sequences of LptF2, LptG2, and LptC2 were aligned with their respective LPS-transporting homologues.

**Supplemental Figure 2: LptC2 does not form a stable interaction with LptF2 or LptG2.** Strep-tagged LptC2 was used in pull down assays to detect protein-protein interactions with FLAG-tagged LptF2 or LptG2. The (*) indicates a non-specific band detected by the anti-strep tag antibody which did not bind to the affinity resin. LptC2 was efficiently purified using the Strep Tactin resin, but neither LptF2 nor LptG2 were detecting in the eluate.

**Supplemental references**

1.	Nierman WC, Feldblyum TV, Laub MT, Paulsen IT, Nelson KE, Eisen JA, Heidelberg JF, Alley MR, Ohta N, Maddock JR, Potocka I, Nelson WC, Newton A, Stephens C, Phadke ND, Ely B, DeBoy RT, Dodson RJ, Durkin AS, Gwinn ML, Haft DH, Kolonay JF, Smit J, Craven MB, Khouri H, Shetty J, Berry K, Utterback T, Tran K, Wolf A, Vamathevan J, Ermolaeva M, White O, Salzberg SL, Venter JC, Shapiro L, Fraser CM. 2001. Complete genome sequence of *Caulobacter crescentus*. Proc Natl Acad Sci U S A 98:4136-41.

2.	Stankeviciute G, Guan Z, Goldfine H, Klein EA. 2019. *Caulobacter crescentus* adapts to phosphate starvation by synthesizing anionic glycoglycerolipids and a novel glycosphingolipid. mBio 10:e00107-19.

3.	Uchendu CG, Guan Z, Klein EA. 2024. Spatial organization of bacterial sphingolipid synthesis enzymes. J Biol Chem 300:107276.

4.	Zik JJ, Yoon SH, Guan Z, Stankeviciute Skidmore G, Gudoor RR, Davies KM, Deutschbauer AM, Goodlett DR, Klein EA, Ryan KR. 2022. *Caulobacter* lipid A is conditionally dispensable in the absence of fur and in the presence of anionic sphingolipids. Cell Rep 39:110888.

5.	Simon R, Priefer U, Puhler A. 1983. A broad host range mobilization system for in vivo genetic engineering: transposon mutagenesis in gram negative bacteria. Bio/Technology 1:784.

6.	Thanbichler M, Iniesta AA, Shapiro L. 2007. A comprehensive set of plasmids for vanillate- and xylose-inducible gene expression in *Caulobacter crescentus*. Nucleic Acids Res 35:e137.
