## Supplemental Figure 1 for "Homologues of the inner-membrane LPS transport proteins are required for sphingolipid transport in *Caulobacter crescentus*"

**Percent identity: 20.85; Percent similarity: 40.05**

```
LptF2 (CCNA 01214)        MSAKDLPNGGPRLIDRYLLRLLLWPLAACLGVTVIALLLERILRLLDVLSQSSARFGYVA 60
LptF (CCNA 01762)         ----------MRLIERYLFRQLLGPTLLATAALVALALLARSLSEFDVLVEQRQSALVFL 50
                                     ***:***:* ** *   . .. *   ** * *  :*** :.      .

LptF2 (CCNA 01214)        QLAANLVPHYLGLALPVAFFVALFIVIVKLSDGSEIDALLASGQSLERIAAPFVAVGVLL 120
LptF (CCNA 01762)         KIILLALPQLLGIMMPLALFVAALVALNRLHTEQEIVVCFAGGMSRWRVISPAMRLAVIA 110
                          ::    :*: **: :*:*:*** ::.: :*   .** . :*.* *  *: :* : :.*:

LptF2 (CCNA 01214)        SVFSIIVFGYMQPYSRYAYRAVMHAAVNAGWNGRLAGGAFIDDKGSLLTADSADI-AGQR 179
LptF (CCNA 01762)         ALISLISGLWLQPWSARQIRETAFQIKTDVAATLVQPGQFTEPG-PGLTVYAQSIDRDNK 169
                          :::*:*   ::**:*    * . .   .      :  * * :     **. : .*  .::

LptF2 (CCNA 01214)        LTRVFIRRLDAKGQEEIITAASADLKMDPDGKTITMDLRDGRRIAHDA---YGTYRNLAF 236
LptF (CCNA 01762)         IQNLFIHQELPNGAASTYSSREAEIATRKGEPVLIMRHGSNQQFSRDGVLQYTSFDEYTF 229
                          : .:**::   :*  .  :: .*::    .  .: *   ..:::::*.   * :: : :*

LptF2 (CCNA 01214)        TSLTTQTPLSAAAVLLRDR--GGDERELTMGELRRE--AKSPDPVVSKGTLLSEFYGRLA 292
LptF (CCNA 01762)         DLS---SLFHS-DELLHYKIADRYPHELFFPDLTQEWEQRNKDKLLAEG--HSRFAGPLY 283
                                : : :   **: :  .   :** : :* :*   :. * ::::*   *.* * *

LptF2 (CCNA 01214)        RAAFLPFLPLLAFPLGLASKRG--NRAPGLIMAGLLLLAFQHSLLLGQGMAKAGKADAFA 350
LptF (CCNA 01762)         NIALMSMA-LAAVLGGSFSRMGYGKRIAAV---GA---AAAIVRIVGFGVQAACEDD--- 333
                          . *:: :  * *.  *  *: *  :*  .:   *    *     ::* *:  * : *

LptF2 (CCNA 01214)        AIWIPFTL---FAALSVWLFVGS---RTRPGDTPVSRLVRRINTLVTHALRLLPQPKKRQ 404
LptF (CCNA 01762)         -PWLNVLQYIVPLAATAWAMSQIFRQRVSPRRK---------------TKPATPAPAFSG 377
                            *: .       * :.* :      *. *  .               :    * *

LptF2 (CCNA 01214)        AA       406
LptF (CCNA 01762)         AA       379
                          **
```

**Percent identity: 23.54; Percent similarity: 44.71**

```
LptG2 (CCNA 01213)       -----MKLQLYVLRTVATRILGAALILMSVLQILDLLDVTSDILDRG-LGMAGVGYYAAL  54
LptG (CCNA 01761)        MMLTMGMLERYVLKRTMGALVGALAVLSAMVMLIAFVDIARNVGTRADASFLRLLYLTIL  60
                                *: ***: .   ::** :*  ::: :: ::*:: ::  *.  .:  : * : *

LptG2 (CCNA 01213)        RLPRLFEQVAPIAVLAGGLFAFSQLARESAIVAMRASGISGYRIVGMAVPAAVAVMLLDA  114
LptG (CCNA 01761)        QAPATILVLAPFIFLFGTMWAFVELNRRSELIAMRAAGISAWRFIMPAAASSFVI---GL  117
                         : *  :  :**: .* * ::** :* *.* ::****:***.:*::  *. ::..:   .

LptG2 (CCNA 01213)        LCGQVLAPRADPTLADWWRNTT--P-VAERKEPVPRTFRAGAD---LVIGANA--SAD-G  165
LptG (CCNA 01761)        LTITLLNPLTTAMTAKFETERDRTMNGYLKEAPKGTWLRQGDDKTQIVIRARARELVDGS  177
                         *   :* * :    *.:  :         ::  *    :* * *   :** *.*   .* .

LptG2 (CCNA 01213)        RTITGVTIF--RRDSKGI--LVEKVEAPAARYDGKAWTLEQPKTTRFA-GDLSQASTAAA  220
LptG (CCNA 01761)        VRLRGVSLFVYTLNAKGVMDFTRRIEANEARLEPGFWRLSGVREATPGAGAIRS----DS  233
                           : **::*    ::**:  :..::**  ** :   * *.  : :  . * : .     :

LptG2 (CCNA 01213)        TSWPTALRPQDVQGLFGDDSM----PTAASARRALENGGSDRPESFYATHLQAAFASPFV  276
LptG (CCNA 01761)        LSIPSNLDDRTASERFNTPQAVALWRLPATIQRTADAGFS---AVPFKLRLQQDLATPLL  290
                          * *: *  : ..  *.  .        *: :*: : * *      :  :**  :*:*::

LptG2 (CCNA 01213)        SLVMLLLSAPVALANFRSGQGAVLLTGGLAAGLMFLVANGMLTALGESGALTPFLAVWAA  336
LptG (CCNA 01761)        FAAMSVLAAAFSLRLMRLGGLAMLAGSGVALGFGFFFFNELCSTLGRADVLAPFVAAWTP  350
                           .* :*:* .:*  :* *  *:*  .*:* *: *:. * : ::**.:..*:**:*.*:

LptG2 (CCNA 01213)        PAIFGALAVRTFLALEG- 353
LptG (CCNA 01761)        PTVALLVGFTLLCYTEDG 368
                         *::   :..  :   *.
```

**Percent identity: 37.10; Percent similarity: 51.58**

```
LptC2 (CCNA_01226)       ---------MLTQSDGAAAILPSQ-TAAERAR-NRRRKPVRRLRLGLAVFAAGVAATVIV  49
LptC (CCNA_03716)        MGGDGHLRPQLKDLMQPAAVMTGRSMSADLARWRRRSRRVRAARLVLPAAIGVLLLAVIA  60
                                   *.:    **:: .:  :*: ** .** : **  ** * .  . :  :**.

LptC2 (CCNA_01226)       QAAWRSLASSKLQTQAATAPLVLDKPRFTGVLKDGRPFLITAERAERDAKDQNIVRLTAP  109
LptC (CCNA_03716)        QVGWRTYLAASRAPAEARTEIRLITPRFYGQSTDGRSFMITARSAIRDDVDPRRIILEEP  120
                         *..**:  ::.     * : : * .*** *  .*** *:***. * **  * . : *  *

LptC2 (CCNA_01226)       LLVRGYGEPNPSQATAKSGVYREAENTLLLTDEVKITSAEGFDFDAPRALIDLRTGAVSG  169
LptC (CCNA_03716)        ALTLDLGSPTPTRMTAKHGVYRQDTFGLNLKDDVRLDDGEGYRFASEESFVDTRTGDVSG  180
                          *. . *.*.*:: *** ****:    * *.*:*:: ..**: * : .:::* *** ***

LptC2 (CCNA_01226)       DAGIAGSGPKGSTRASAYEVTDKGDRVVLKGGVRTRLEPRP    210
LptC (CCNA_03716)        ESTMNGEGPSGQVQSSAYSVYDKGDRIVFRGGVRARFEQQ-    220
                         :: : *.**.*..::***.* *****:*::****:*:* :
```
