## Supplementary figures and images for "Homologues of the inner-membrane LPS transport proteins are required for sphingolipid transport in *Caulobacter crescentus*"

### Supplemental Figure 2

Supplemental Figure 2

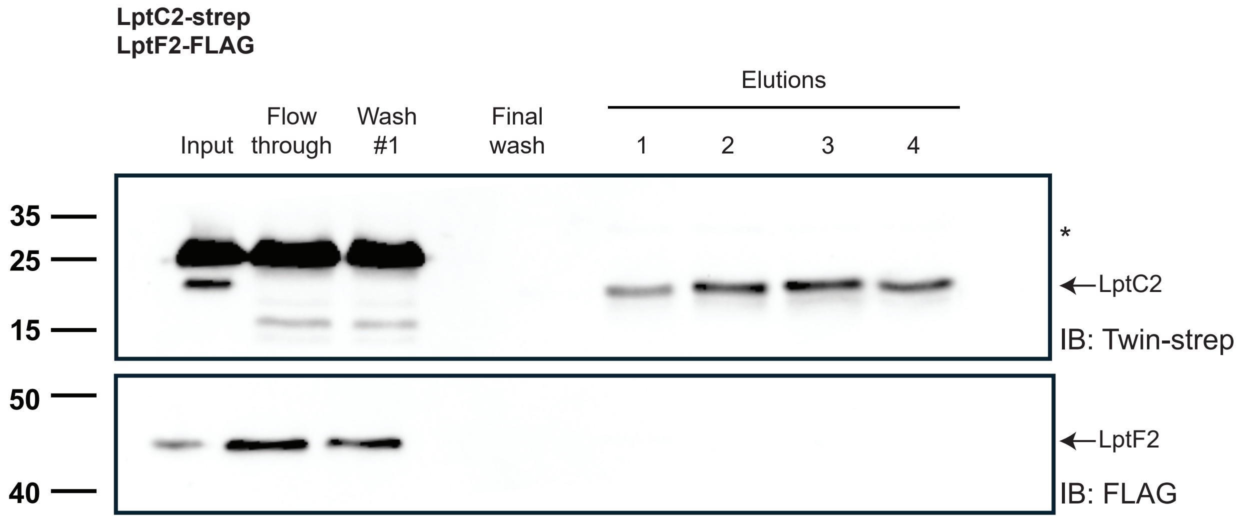


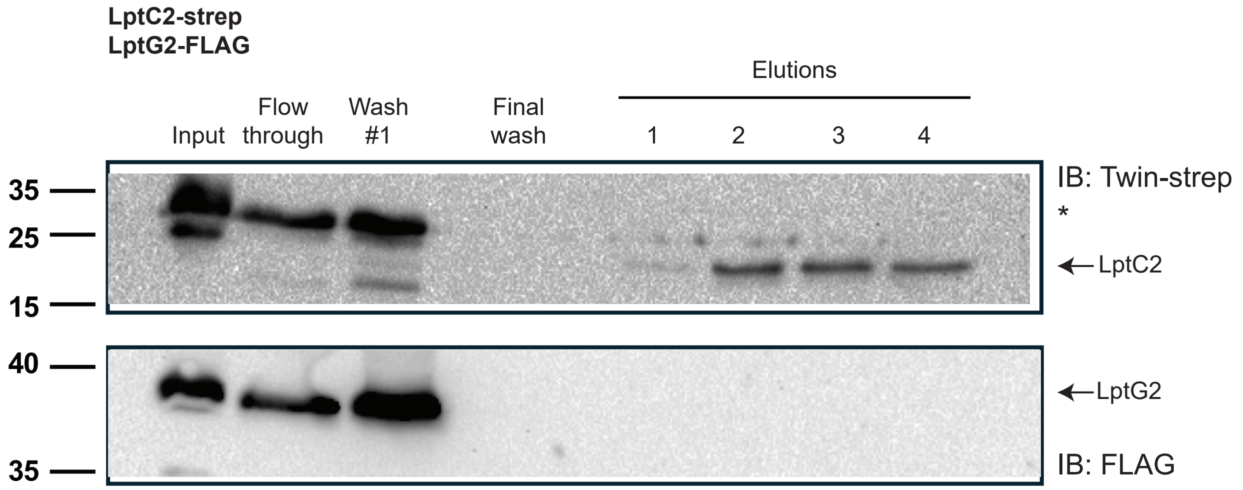
